# Comparing the impact of contextual associations and statistical regularities in visual search and attention orienting

**DOI:** 10.1101/2024.04.12.589295

**Authors:** Marcus Sefranek, Nahid Zokaei, Dejan Draschkow, Anna C. Nobre

## Abstract

During visual search, we quickly learn to attend to an object’s likely location. Research has shown that this process can be guided by learning target locations based on consistent spatial contextual associations or statistical regularities. Here, we tested how these different types of learning aid the utilisation of established memories for different purposes. Participants learned contextual associations or statistical regularities that predicted target locations within different scenes. The consequences of this learning for subsequent performance were then evaluated on attention-orienting and memory-recall tasks. Participants demonstrated facilitated attention-orienting and recall performance based on both contextual associations and statistical regularities. Contextual associations facilitated attention orienting with a different time course compared to statistical regularities. Benefits to memory-recall performance depended on the alignment between the learned association or regularity and the recall demands. The distinct patterns of behavioural facilitation by contextual associations and statistical regularities show how different forms of long-term memory may influence neural information processing through different modulatory mechanisms.

## Introduction

Memories may be about the past, but we use them proactively to anticipate events about to unfold [1]. For example, when searching for a restaurant we have visited in the past, relevant information in long-term memory can guide our search. Long-term memories supporting visual search come in different flavours and are supported by different brain systems [2–5]. For example, we might have learned that most restaurants within a given city are on one side of the main road. This *statistical* regularity can guide our attention to that side of the road when searching for a restaurant [6,7]. Learning of such statistical regularities develops gradually and is supported by striato-cortical learning mechanisms [8–11]. In contrast, we may form a specific memory association between the location of a particular restaurant and its *contextual* surroundings. These contextual associations can guide our attention to the specific location of the remembered restaurant. Contextual associative memories form quickly and depend on plasticity mechanisms involving the medial-temporal lobe [12–14].

Visual-search studies have shown how both types of long-term memory can guide attention to improve performance. Studies that manipulate probabilistic or statistical regularities of the location of the target within an array of distractors show strong and lasting performance benefits for targets that appear at more frequent locations [15]. Studies that manipulate contextual associations show improved performance for targets that appear in a consistent location within a specific configuration of distracting items [13,16,17]. How statistical and contextual memories guide attention and performance has mainly been studied separately.

Attentional guidance and behavioural facilitation by statistical regularities has been shown primarily by manipulating probabilistic learning [15,18–21]. In probabilistic learning tasks, participants learn that a target occurs more frequently within a consistent region of the display, such as a specific half [22], quadrant [7,23], or hotspot [24]. This generalisable probabilistic information can facilitate visual search as a “probability cueing” effect. Such learned environmental statistics can also be used to guide attention [25]. Probability cueing has been characterized as “habitual” in that the attentional bias persists even when the cue is devalued [23,26] or when the task switches (25, e.g., from a letter search to a scene search task).

Attentional guidance by contextual associations has mainly been studied by "contextual cueing” manipulations in visual-search tasks using simple stimulus arrays [16,27–29]. In these tasks, the target location varies across the entire display over trials, but some specific repeating array configurations predict the precise target location. These repeating contexts facilitate visual search via reduced reaction times. The benefits of contextual memory associations have also been demonstrated when searching for consistently located targets within repeated scenes [30–35] and immersive virtual and real environments [36,37]. The benefits of contextual cueing extend to memory recall of target identity and location in healthy and at-risk populations [38–44]. Also, studies using learned scenes as attention-guiding memory cues for subsequent target identification have revealed memory-guided signals that proactively prepare anticipatory attention to anticipate target locations [45–49].

Critically, statistical regularities and contextual associations may work together to guide behaviour [5,11,50–52]. However, due to differences in methodologies used to study the two types of memories, direct comparisons of their behavioural consequences are rare. Notably, Goldfarb et al. [53] measured visual-search performance for targets abiding by either probabilistic or contextual regularities. Both types of association yielded significant learning, as measured by decreasing response times over the course of the visual-search task. Concurrent brain-imaging measures suggested the effects were supported by distinct neural systems. Trial-evoked blood oxygenation level dependent (BOLD) responses in the hippocampus predicted response-time benefits in contextual-association trials, while BOLD in the striatum predicted response-time benefits in probabilistic-association trials. This important study elucidated that contextual and probabilistic regularities guide visual search through dissociable neural systems and opened additional interesting questions. Do similar patterns of search benefits generalise to visual search within natural scenes? Are there differences in how these two types of memory guide attention and, consequently, impact perception? Do they impact subsequent memory-related decisions in different ways? We sought to explore these questions.

In our study, we compared how statistical regularities and contextual associations guide learning to improve visual search performance. After effective learning, we probed how these two types of memories modulated attention and memory functions. We manipulated statistical regularities and contextual associations for the location of target stimuli appearing within natural scenes. We chose natural scenes to approximate everyday search behaviour [30–32,54]. By using different scene categories, we manipulated the type of association that could support learning of target locations within the same visual-search task. For two scene categories, the target location followed a deterministic statistical regularity, consistently appearing within the same side of the screen depending on the category (left or right). For two other scene categories, target locations were specifically tied to their context within a given scene. The target could vary between left or right for the scene category, but it always appeared within the same location in the same scene. For the two remaining scene categories, the target location varied randomly and was not predictable according to scene category or scene identity. Participants were not informed about the statistical regularities and contextual associations depending on scene category. Individual scenes from the three conditions – contextual, statistical, and random – were randomly intermixed within each learning block, enabling the comparison of different types of learning benefits without state-related confounds. After completing multiple blocks of the visual-search task, participants completed a memory-guided attention-orienting and a memory recall task to compare the impact of different types of memories.

We hypothesized that statistical regularities and contextual associations would have different time courses for benefitting visual-search performance, with statistical learning improving search more slowly than contextual associations (see also 53, Fig S1). We used different scene-to-target intervals in the memory-guided attention-orienting task to ask whether statistical regularities and contextual associations guided attention with different time courses. Following the attention-orienting task, participants completed a memory recall task. We hypothesized that performance in this explicit memory task would tap into the nature of the association learned incidentally during visual search. Memory for the target side should be aided by previously learned lateralised statistical regularity, whereas memory for the precise target location should be aided by previously learned contextual associations.

## Methods

The study was approved by the Central University Research Ethics Committee of the University of Oxford code: R63062/RE001.

### Participants

Participants were recruited online using Prolific (www.prolific.co), an online crowdsourcing platform. Recruitment began on 14/12/2021 and finished on 13/01/2024. Participants were fluent in English, aged 18-40, and reported having no cognitive impairment, dementia, or history of mental health conditions. Participants were chosen only if they had successfully completed more than 10 studies previously on the platform and had an approval rating of over 90% on prolific. They were compensated £10 per hour for their participation.

The full study included three sessions over three separate days (see below). Progressing to the next day required a minimum level of performance. Based on a related pilot study in lab, a power analysis in G*Power [55] targeted the detection of medium effects in the memory-guided attention-orienting task (d= 0.4, α = 0.05,1-β = 0.95) suggested a sample size of n = 84. To yield this final number of participants, the initial sample recruited consisted of 170 individuals. Participants who performed at near-chance level (in any of the task conditions) were excluded as a precaution to avoid using data from online participants who were inattentive or could not/would not perform at a baseline that was expected based on prior lab-based experiments [56]. After Day 1, 38 participants were excluded (29 based on performance and 9 based on technical malfunction). The remaining 132 participants were invited to the session on Day 2. Of these, 20 did not complete the task, and an additional 8 were excluded (7 based on performance and 1 based on technical malfunction). The remaining 104 participants were invited to the final session on Day 3. Seven did not complete the task, and 13 were excluded from the analyses (11 based on performance and 2 based on technical malfunction). Technical malfunctions entailed problems saving the experimental data or re-routing participants correctly between segments of the study. The final sample consisted of 84 participants who completed all sessions to a satisfactory level (age range: 21 to 40 years; mean age: 29 years; 42 females, 78 right-handed). Demographic information was only collected for participants who made it to the final day of the experiment.

### Apparatus

The experimental task was programmed using Psychopy [57] and hosted on the online platform Pavlovia (https://pavlovia.org/). Participants completed the experiment on their own personal computer. They were instructed to sit one arm’s length away from their computer screen and to adjust the screen brightness to ensure good visibility.

### Stimuli

One hundred and sixty-eight natural scenes were used in the experiment. Scenes were drawn from six different categories in equal numbers (28 bathroom, beach, desert, forest, gym, and kitchen scenes). Of these, 84 unique scenes were randomly chosen to be included in the experiment for each participant (14 from each category). Four additional scenes that were not from any of these categories were used for the practice trials. The size of each scene was 768 x 768 pixels. Scenes were displayed centrally with a grey background filling the rest of the internet browser window.

Semi-transparent, multi-coloured images of a bell and a dog were used as target stimuli. Participants searched for either of these two stimuli embedded within each scene. Target location was determined according to three experimental memory-association conditions (Contextual, Statistical, or Random). The target image was 23.04 x 23.04 pixels and appeared randomly rotated on each trial.

### Procedure

A shortened demo version of the learning task can be completed at (https://run.pavlovia.org/msefranek/hipstr_practice). Participants were provided with a link by Prolific (www.prolific.co), which rerouted them to the experiment on Pavlovia (pavlovia.org). After reading through a description of the study, participants provided informed consent. Next, they completed a set of 4 practise trials to become accustomed to the procedure.

The experiment consisted of three sessions. The first two sessions made up the learning phase (visual search). On the third day, participants completed a surprise memory-guided attention-orienting task and, after a pause, a surprise memory recall task.

### Visual search

During visual search, participants searched through each of 84 scenes for the target (bell or dog) and made a speeded discrimination response as soon as they found the target. Target identity was randomised, and participants made a two-alternative forced choice to indicate if the target was the bell or the dog. In each session (Day 1, Day 2), they completed five blocks of searching through the 84 scenes. Each learning block took roughly 6 minutes to complete resulting in the entire learning session being about 30 minutes in duration. The order of the scenes was randomised. After each block, participants had the opportunity to take a short break and to continue when they felt ready.

Fig 1a shows the experimental procedure for a given learning trial. First, a grey screen with a white fixation cross appeared for 500 ms. This was followed by a brief (250 ms) preview of the scene alone, with no embedded target. This served as a prompt for the upcoming scene with the embedded target. After a 750 ms interval with a grey background, the scene was presented again, this time with an embedded target. Participants had up to 7 seconds to search for the target in the first block and respond with the correct keystroke (“D” key corresponding to the dog target and “L” corresponding to the bell). For the remaining blocks, search time was reduced to 5 seconds. If participants responded before the allotted time elapsed, the search window closed. After responding, centrally presented feedback for 500 ms indicated if their response was “Correct” or “Incorrect”. If the participant was unable to find the target within the allotted search time, “Incorrect” was presented as feedback. Trials were separated by an inter-trial interval (ITI) of 200 ms.

**Fig 1.**
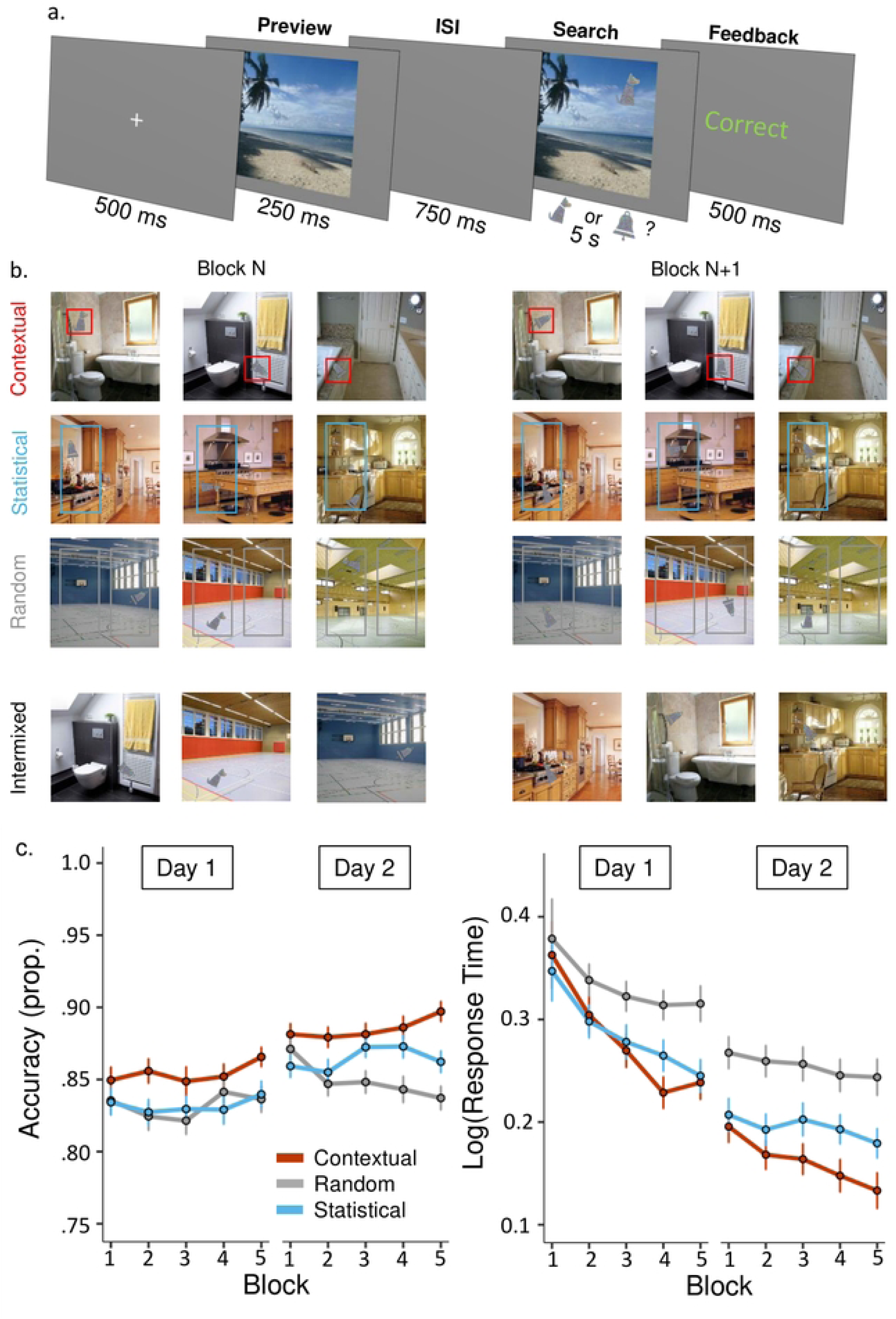
Learning task and performance plots. (a) A schematic of an individual trial in the visual-search task. (b) A diagram illustrating the different types of learning associations across blocks. For scene categories in the ‘Contextual’ condition, the target appeared in a consistent location relative to the scene identity. Across scenes in this category, targets could appear on the left or right but, within the same scene, the target always appeared in the same location, here highlighted in red. For scene categories in the ‘Statistical’ condition, targets appeared on a consistent side according to the scene category, here highlighted in blue. The specific location of the target within the consistent side could vary according to scene identity. For scene categories in the ‘Random’ condition, targets could appear at any location within the left or right side of the scene, with no consistency across blocks, here highlighted in grey. The bottom, ‘Intermixed’, panel illustrates that all three conditions were randomly intermixed within any given experimental block. (c) Accuracy (left graph) and Response Times (right graph) plots for each of the conditions over the 5 learning blocks in each of the visual-search sessions. Response times were log-transformed to approximate the normal distribution. Overall, participants demonstrated significant benefits from Contextual associations (accuracy and response times) and Statistical regularities (only response times) compared to the Random condition. Learning performance also improved from Day 1 to Day 2. Error bars were calculated using the “summarySEwithin” function (Version 1.5.1).

Target locations during the visual-search phase (Fig 1A) were determined according to three experimental conditions: Contextual, Statistical, or Random. The six categories were split evenly into these 3 conditions, with 2 scene categories per condition, and 14 unique scenes in each category (3 conditions x 2 scene categories x 14 unique scenes = 84 total scenes). For the two scene categories with Contextual associations, the location of the target remained the same for a given scene across all five blocks. In other words, there was an exact match between a scene and the target location within it. For example, if the target appeared on the window of a house in a scene in block one, the target always appeared in the same location on the window in that scene in the subsequent four blocks. Independent of scene category, for half the scenes this exact target location was on the left hemifield and for the other half it was on the right hemifield. For the two scene categories with Statistical regularities, the target consistently appeared in the same hemifield across the five blocks, but the exact location in that hemifield varied randomly. For example, for one participant, targets would always appear on the left for gym scenes and on the right for beach scenes. Finally, for the two scene categories with Random associations, the target location varied randomly and could not be predicted by scene category or identity.

The assignment of conditions to scene categories was counterbalanced across participants. Target locations could only occur within a specific region of pixel space defined within Psychopy. On the x-axis, possible locations were in the central 50% of the right or left hemifield (.25 to .75, if 0 is the middle and 1 is the edge of the scene). On the y-axis, possible locations ranged from the midpoint to 75% of the possible space in the upper or lower hemifield (0 to .75). The same boundaries for target placement were used for all experimental conditions. For scenes in the Contextual condition, the initial location was drawn randomly, and this specific location was preserved for each subsequent learning block. For scenes in the Statistical condition, coordinates in each block were drawn randomly from values within the hemifield tied to the scene category. For scenes in the Random condition, a new draw would determine the target location in each block.

### Memory-guided attention

On Day 3, participants completed a surprise memory-guided attention-orienting task (Fig 2a). The same scenes were used as in the Learning Phase, but they were presented much more quickly. The order of the scenes was randomised. In each trial, after a brief preview of the scene alone, which acted as a memory cue, participants discriminated whether the bell or dog target was embedded in the scene when briefly re-presented.

**Fig 2.**
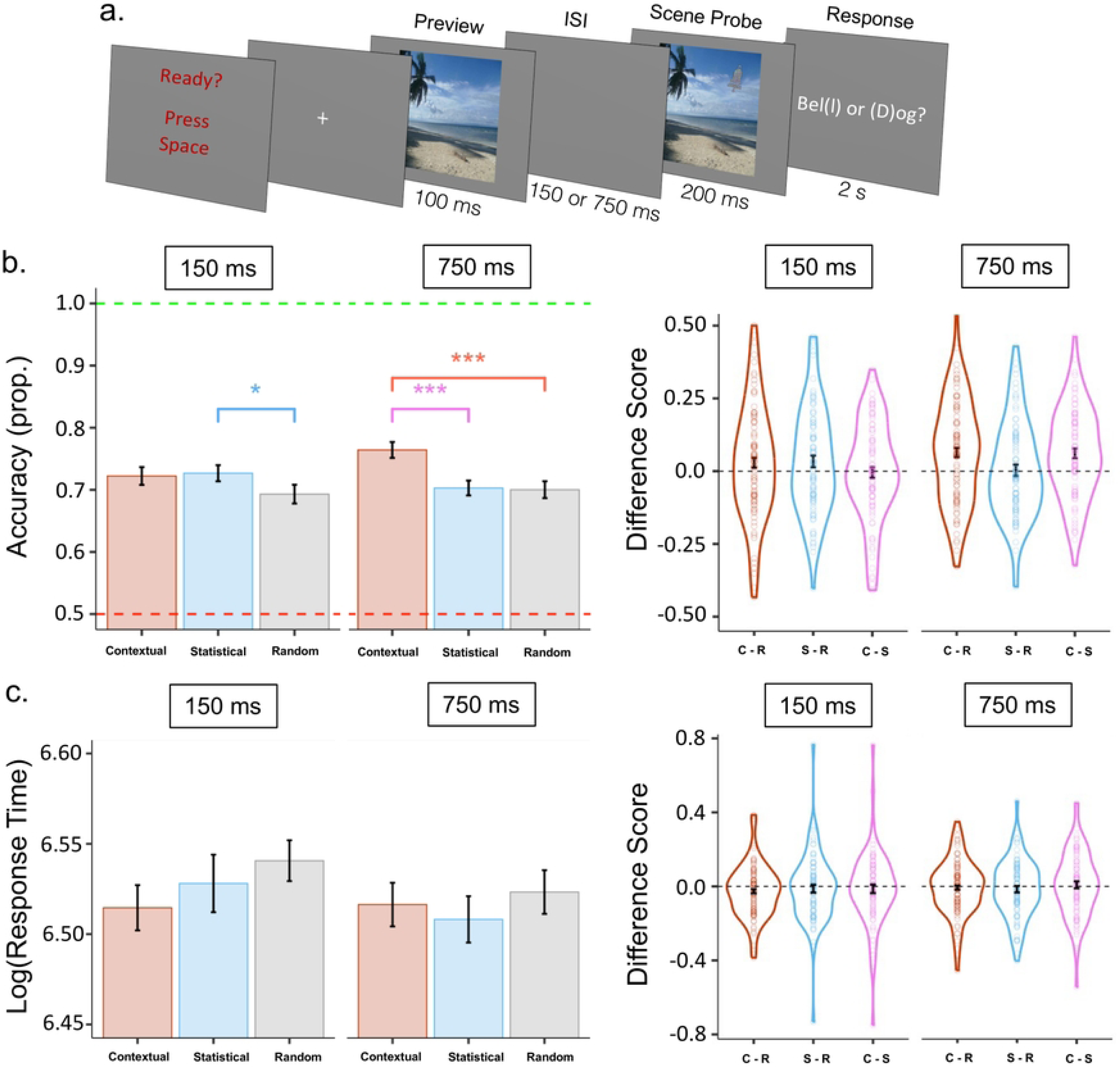
Memory-guided attention task and results. (a) A schematic of the attention-orienting task. (b) On the left, response accuracy is plotted separately based on shorter (150-ms) or longer (750-ms) ISI. At the shorter ISI, accuracy was marginally higher only for Statistical associations relative to the random condition. After the longer ISI, accuracy was higher for Contextual associations relative to both other conditions. Additionally, within each participant, accuracy was compared between conditions to generate the Difference Score plotted on the right. Each dot corresponds to the Difference Score of a single participant in a given comparison of association conditions (e.g., C-R represents Contextual association accuracy minus Random association accuracy). (c) There were no significant differences in response times based on the type of learned association or ISI. The Difference Score for response time was calculated in the same manner as for accuracy. Error bars specify normalized mean standard error.

So that participants could prepare for the quick pace of the ensuing trial, each trial began with a screen that stated “Ready? Press space.” After a subsequent fixation cross (750 ms), one of the scenes flashed for 100 ms with no embedded target. After a short (150 ms) or long (750 ms) blank inter-stimulus interval, the scene flashed again for 200 ms, this time containing the target. A grey background then appeared with the question “Bell or Dog?” in white font. Participants had 2 s to report the target they saw using the “D” key for dog and the “L” key for bell. After participants responded to the target identity or ran out of time, the next trial immediately began. Target location matched the rules from the learning task. In each Contextual trial, the target appeared in the exact same location as in the learning phase. In Statistical trials the target appeared in the same hemifield as in the learning phase. In Random trials the location was random.

### Memory-recall

On Day 3, after completing the memory-guided attention task, participants completed an unrelated demographics questionnaire, which lasted 8.2 minutes on average. This was followed by a surprise memory-recall task (Fig 3a). Participants were tested on their location memory for the target-scene associations they had learned throughout the experiment. Specifically, they were instructed to “Click the exact location you remember seeing the target during the previous two days of the study.” They were not given specific instructions on which locations to recall on scenes in the Statistical and Random conditions to keep them blinded to the experimental manipulations.

**Fig 3.**
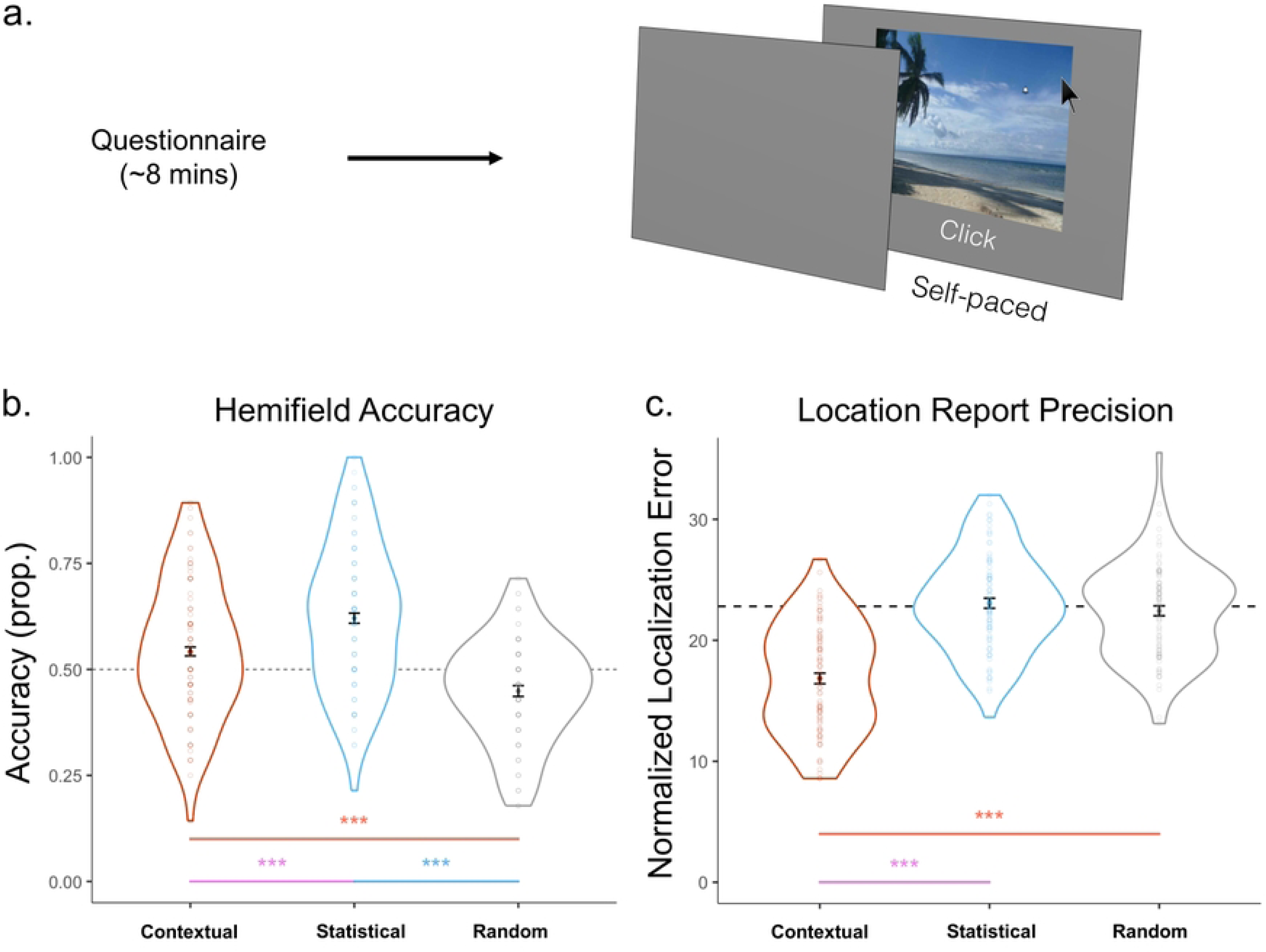
Memory-recall task and performance plots. (a) A schematic of the recall task. If participants recalled (clicked) in the same hemifield as the orienting target, this counted as a correct location accuracy report for the proportion in (b). The distance between the location participants recalled and the orienting target location represented their report precision (c). (b) Hemifield accuracy was significantly higher in the Statistical and Contextual associations than in the Random associations, and significantly higher in the Statistical than in the Contextual condition. (c) Location error results. Only trials in which participants placed the target in the correct hemifield are included. Location error was significantly lower in the Contextual than in the Statistical and Random association conditions. The Statistical and Random conditions did not differ from chance. Error bars represent normalized mean standard error. Dotted lines represent chance value.

In each trial, participants viewed each of the scenes at their own pace and used the mouse to click the location where they remembered seeing the target. Trials were presented in random order. After responding, an inter-trial interval of 750 ms occurred until the next scene appeared.

### Data analysis

Parametric analyses were applied, transforming the data when necessary to conform to the required assumptions. When main effects were significant, planned paired samples t-tests were applied to compare levels of the experimental variables (i.e., between different learned associations). In line with our a-priori hypotheses, throughout the visual-search, attention-orienting, and memory-recall results, t-tests comparing learning conditions (Contextual or Statistical) and the Random condition were one-sided. Instead, t-tests comparing Contextual and Statistical effects were two-sided. Comparisons between effects on Day 1 versus Day 2 of learning were two-sided. Comparisons within attention-orienting ISI (e.g., contextual condition at 150 ms vs 750 ms ISI) were two-sided. The Greenhouse-Geisser correction was applied when the sphericity assumption was violated. Throughout the experiment, to account for within-subject comparisons (i.e., between association conditions), we followed procedures from Morey [58]. Specifically, we calculated error bars using the “summarySEwithin” function (Version 1.5.1), which normalizes data to remove between-subject variability and then calculates the variance from this normalized data.

All analyses were conducted using RStudio [59]. Descriptive statistics and paired sample t-tests were conducted using the “stats” package in R [60]. Within-subject ANOVAs were conducted using the “ez” package when necessary to aggregate data (61;Version 4.4). All graphs were produced using the “ggplot2” package [62].

### Learning

To analyse performance on the learning task, we tracked target identification accuracy (% of correctly identified targets) and response time over the 5 blocks for each day. The independent variables of interest were Association (Contextual, Statistical, Random), Block (1 to 5), and Day (1 or 2). We only examined response times for trials in which participants correctly identified the target. Response-time outliers (above 2 standard deviations from the individual mean across trials of all conditions) were removed from analysis (5.5% of trials on Day 1 and 5.7% on Day 2). The Box-Cox procedure [63] was used to test the distribution and residuals of response-time values. Based on this, we applied a log transformation to response times on both days of learning to approximate the normal distribution and to meet ANOVA assumptions.

### Memory-guided attention

We assessed memory-guided attention by examining target identification accuracy (% of correctly identified targets) and response times. The independent variables of interest were memory association (Contextual, Statistical, Random) and ISI (150 ms or 750 ms; Figs 2b and 2c). Only trials in which the target was correctly identified in the attention task were used to calculate the response-time averages. Also, response-time outliers (above 2 standard deviations from the individual mean across trials of all conditions) were removed from the analysis (4.2% of trials). The Box-Cox procedure [63] recommended log transformation for the response-time values.

### Memory-recall

To assess performance on the surprise memory-recall test, we calculated participants’ accuracy at locating the target in the correct hemifield as well as their precision in placing the target at a learned location (Figs 3b and 3c). The independent variable of interest was memory association (Contextual, Statistical, Random).

Hemifield accuracy was assessed by determining the proportion of trials in which participants correctly localised within the hemifield in which the target occurred during the orienting task. To elaborate, during the orienting task, the target flashed once for any given scene. During memory recall, if participants clicked in the hemifield where this target occurred during the orienting task, we determined that they responded accurately (Fig 3b). Then, for these correct-hemifield trials only, location precision was calculated by measuring the distance in two-dimensional space between the centre of the target location in the orienting task and the participants’ response during the memory-recall task (Fig 3c). We focused on these correct hemifield trials to demonstrate that, beyond responding to the target’s hemifield, precision in the Statistical and Random conditions was at chance (Fig 3c). In contrast, participants were able to localise the target location, beyond just its side, in the Contextual condition. To simplify error values from the raw Psychopy output coordinates, error values were divided by the maximum localisation error, which generated a normalised percentage error out of 100.

A specific hemifield memory chance error value was calculated for each individual participant, for correct identification report trials only (this was used in Fig 3c). Response locations in the left hemifield and in the right hemifield for the corresponding scenes were shuffled with responses to other scenes with left and right response locations (e.g., left hemifield target locations shuffled with other left hemifield target locations). The localisation error was then recalculated using these shuffled values for each participant over 1000 iterations. These new localisation error values were then averaged to generate the chance-performance value.

In both hemifield accuracy and location precision analyses, we measured participants’ memory recall performance compared to the orienting target rather than the learning target locations. One of the important metrics we analysed was memory recall accuracy (Fig 3b). This result examined the proportion of memory recall trials in which participants placed the target in the same hemifield as it occurred during the orienting task. This calculation could only be made using the orienting target location. For the Random condition, locations during the learning task varied between hemifields, meaning that there was no intuitive hemifield to use to calculate accuracy. Instead, the hemifield where the random target occurred during the orienting task provided a natural anchor point. Therefore, using the orienting target location as the comparison point allowed us to calculate this important result.

## Data availability

Raw data, pre-processing scripts, pre-processed data, and final analysis scripts can be found on: OneDrive (upon acceptance, upload to GitHub will follow).

## Results

In an online experiment conducted over three consecutive days, we investigated how contextual associations vs. statistical regularities about the locations of target objects in natural scenes changed performance (relative to a random-association baseline) on visual search and subsequent memory-based attention orienting and memory recall.

### Reliable contextual and statistical cueing during visual search

Fig 1a shows the visual-search task containing the different types of memory associations (Contextual, Statistical, Random) and performance in the two visual-search sessions.

An ANOVA showed that accuracy in the visual-search task was significantly influenced by the type of memory and session day (see Table 1). Mean accuracy was 87% (*SD* = .09) in the Contextual condition, 85% (*SD* = .10) in the Statistical condition, and 84% (*SD* = .09) in the Random condition. The memory effect showed that participants made a higher proportion of correct target discriminations in the Contextual relative to the Random condition (*t*(83) = 3.221, *p* < .001, Cohen’s *d* = .441) and Contextual relative to the Statistical condition (*t*(83) = 2.247, *p* = .027, Cohen’s *d* = .303). Contrary to our predictions, there was no performance benefit in the Statistical condition relative to the Random condition (*t*(83) = .926, *p* = .179, Cohen’s *d* = .114). Accuracy was better on Day 2 (mean = 87%, *SD* = .09) than Day 1 (mean = 84%, *SD* = .10) learning (*t*(83) = 6.283, *p* < .001, Cohen’s *d* = .511). There was no effect of Block or any interaction between the variables.

**Table 1.**
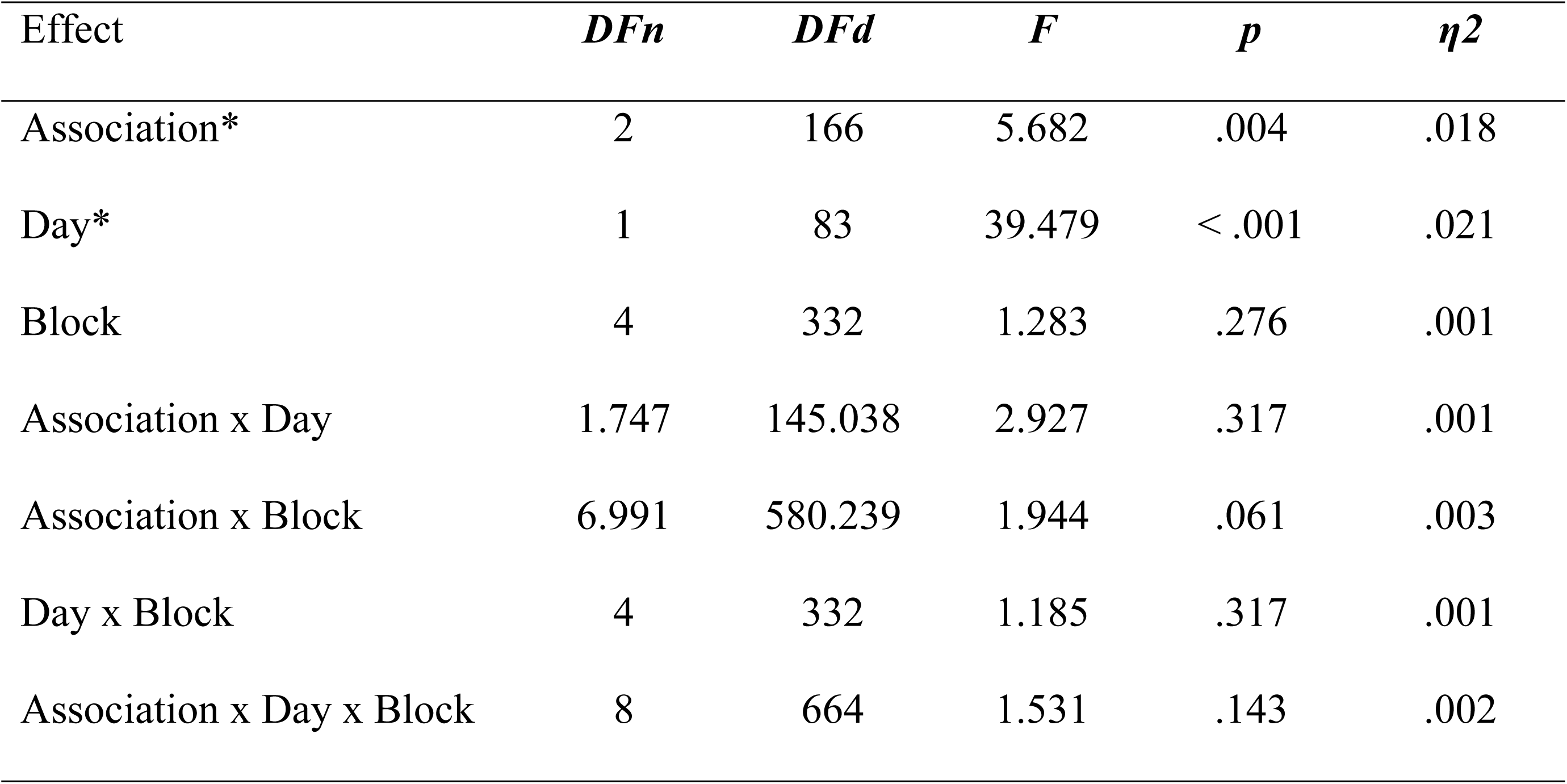
Analysis of accuracy for days 1 and 2 of visual search. There were significant main effects of Association and Day. An asterisk (*) signifies a p-value below .05.

| Effect | *DFn* | *DFd* | *F* | *p* | *η*<sup>2</sup> |
| --- | --- | --- | --- | --- | --- |
| Association* | 2 | 166 | 5.682 | .004 | .018 |
| Day* | 1 | 83 | 39.479 | < .001 | .021 |
| Block | 4 | 332 | 1.283 | .276 | .001 |
| Association x Day | 1.747 | 145.038 | 2.927 | .317 | .001 |
| Association x Block | 6.991 | 580.239 | 1.944 | .061 | .003 |
| Day x Block | 4 | 332 | 1.185 | .317 | .001 |
| Association x Day x Block | 8 | 664 | 1.531 | .143 | .002 |

An ANOVA based on response times during visual search revealed significant effects according to Association, Day, and Block. In addition, the Association factor interacted with both Day and Block, separately. No other interactions were significant (see Table 2). Analyses are shown for log-transformed response times, which were used to approximate the normal distribution.

**Table 2.** Analysis of response time for days 1 and 2 of learning. There were significant main effects of Association, Block, and Day. Interactions occurred between Association x Day and Association x Block. Analysis is shown for log-transformed RT values. An asterisk (*) signifies a p-value below .05.

| Effect | *DFn* | *DFd* | *F* | *p* | *η*<sup>2</sup> |
| --- | --- | --- | --- | --- | --- |
| Association* | 2 | 166.00 | 6.715 | .002 | .023 |
| Day* | 1 | 83.00 | 70.939 | < .001 | .049 |
| Block* | 1.634 | 135.583 | 10.005 | < .001 | .050 |
| Association x Day* | 2 | 166.00 | 7.129 | < .001 | .002 |
| Association x Block* | 6.871 | 570.333 | 3.049 | .003 | .002 |
| Day x Block | 1.618 | 134.256 | 2.853 | .070 | .004 |
| Association x Day x Block | 6.834 | 567.219 | 0.741 | .634 | .001 |

The main effect of Association demonstrated faster response times overall in both Contextual and Statistical associations relative to Random Associations (Fig 1c) (Contextual x Random: *t*(83) = 3.657, *p* <.001, Cohen’s *d* = .474; Statistical x Random: *t*(83) = 2.972, *p* =.002, Cohen’s *d* = .372). Overall, the response times in the Contextual and Statistical conditions did not differ significantly (*t*(83) = .918, *p* =.361, Cohen’s *d* = .137). The main effects of Day and Block indicated that response times decreased as the Days and Blocks of testing progressed. Response times improved over Days for all three conditions (Contextual: *t*(83) = 9.923, *p* <.001, Cohen’s *d* = .702; Statistical: *t*(83) = 8.385, *p* <.001, Cohen’s *d* = .554; Random: *t*(83) = 6.211, *p* <.001, Cohen’s *d* = .447). Response times also decreased over blocks across the days for all three conditions, as indicated by comparing response times in the first vs. last block of the visual-search task across the sessions (Contextual: *t*(83) = 5.75, *p* <.001, Cohen’s *d* = .481; Statistical: *t*(83) = 4.781, *p* <.001, Cohen’s *d* = .407; Random: *t*(83) = 2.397, *p* =.019, Cohen’s *d* = .229).

The interaction between Association and Day likely indicates that the two memory conditions differed more from the Random condition than from one another on Day 1 and differed more from one another on Day 2. On Day 1, mean of log(RT) was .28 (*SD* = .23) in the Contextual condition; .29 (*SD* = .20) in the Statistical condition, and .33 (*SD* = .25) in the Random condition. On Day 2, the mean of log(RT) was .16 (*SD* = .17) in the Contextual condition; .19 (*SD* = .16) in the Statistical condition, and .25 (*SD* = .18) in the Random condition. On Day 1, response times were faster for both Contextual and Statistical conditions compared to the Random condition (Contextual x Random: (*t*(83) = 2.447, *p* = .008, Cohen’s *d* = .312; Statistical x Random: (*t*(83) = 2.473, *p* =.008, Cohen’s *d* = .300) but the two memory conditions did not differ from one another (*t*(83) = .238, *p* = .813, Cohen’s *d* = .034). A similar pattern occurred on Day 2. Response times were faster for both Contextual and Statistical conditions relative to the Random condition (Contextual x Random: *t*(83) = 4.526, *p* < .001, Cohen’s *d* = .588; Statistical x Random: (*t*(83) = 3.202, *p* < .001, Cohen’s *d* = .395) but the difference between memory conditions was not significant (Contextual x Statistical: (*t*(83) = 1.555, *p* = .124, Cohen’s *d* = .223). The three-way interaction (Association x Day x Block) was also not significant, suggesting a similar pattern of improvements over blocks over the two days.

The interaction between Association and Blocks indicated stronger response time gains over blocks for the Contextual condition relative to the Statistical and Random conditions. Across days, the difference between response times in the first vs. the last block in each session was higher in the Contextual condition relative to the Random condition (*t*(83) = 3.057, *p* < .001, Cohen’s *d* = .229) and relative to the Statistical condition (*t*(83) = 2.138, *p* = .036, Cohen’s *d* = .125). The changes in response times between the Statistical and Random conditions did not reach significance (*t*(83) = 1.54, *p* = .063, Cohen’s *d* = .110).

### Effects of contextual and statistical associations on attention-orienting

Contextual and Statistical memories benefitted accuracy measures during the memory-guided attention-orienting task differently depending on the interval between the memory cue and the subsequent target array requiring a perceptual discrimination response. The two-way ANOVA revealed a main effect of Association (*F*(2, 166) = 5.421, *p* = .005, *η2* = .017, Fig 2b) and an interaction between Association and ISI (*F*(1.783, 147.971) = 3.349, *p* = .043, *η2* = .009). There was no main effect of ISI (*F*(1, 83) = .540, *p* = .464, *η2* = .001).

To understand the pattern of effect in the omnibus ANOVA, we compared the effects of Association for the short and long ISIs separately. At the short ISI, mean accuracy was 72%, (*SD* = .14) in the Contextual condition; 73% (*SD* = .14) in the Statistical condition; and 69%, (*SD* = .16) in the Random condition. There was only a one-tailed significant difference between the Statistical and Random conditions (*t*(83) = 1.693, *p* = .047, Cohen’s *d* = .226). There was no difference between the Contextual and Random (*t*(83) = 1.353, *p* = .090, Cohen’s *d* = .193) or the Contextual and Statistical conditions (*t*(83) = .243, *p* = .809, Cohen’s *d* = .032). In the long-ISI conditions, mean accuracy was 76% (*SD* = .12) in the Contextual condition, 70% (*SD* = .14) in the Statistical condition, and 70% (*SD* = .16) in the Random condition. Participants were significantly more accurate for the Contextual condition compared to both Statistical (*t*(83) = 3.620, *p* < .001, Cohen’s *d* = .474) and Random (*t*(83) = 3.330, *p* < .001, Cohen’s *d* = .447) conditions. There was no different in accuracy between Statistical associations and Random (*t*(83) = .151, *p* = .440, Cohen’s *d* = .018).

The equivalent ANOVA comparing response times yielded no significant effects (Fig 2c). Prior to analysis, response time data were multiplied by 1000 (to keep values positive) then log-transformed to approximate the normal distribution and meet ANOVA assumptions. The means for log of RT were 6.52 (*SD* = .20) in the Contextual condition, 6.52 (*SD* = .22) in the Probabilistic condition, and 6.53 (*SD* = .20) in the Random condition. There was no main effect of Association: *F*(2, 166) = .749, *p* = .474, *η2* = .001, ISI: *F*(1, 83) = 1.380, *p* = .243, *η2* = .001, and no interaction: *F*(1.729, 143.493) = .546, *p* = .551, *η2* < .001.

### Contextual and statistical associations can be reliably recalled from memory

The final phase of the experiment was an explicit recall task in which participants viewed scenes from the previous session and reported the remembered location of the target by clicking the mouse cursor within the scene.

To test whether Statistical regularities resulted in explicit memory for target side, we determined the proportion of trials in which the memory recall response (Fig 3a) was placed in the same hemifield as the orienting target. The side and location of the Random target was measured relative to the location in the attention-orienting task in order to provide a consistent anchor (see Methods). Mean hemifield accuracy was 54% (*SD* = .16) in the Contextual condition, 62% (*SD* = .17) in the Statistical condition, and 45% (*SD* = .12) in the Random condition. The ANOVA on hemifield accuracy yielded significant effects based on Association type (*F*(2, 166) = 53.822, *p* <.001, *η2* = .176; Fig 3b). Planned paired-samples t-tests showed that participants’ hemifield accuracy was higher in Statistical (*t*(83) = 9.432, *p* < .001, Cohen’s *d* = 1.157) and Contextual (*t*(83) = 5.710, *p* < .001, Cohen’s *d* = .654) associations compared to the Random condition. Furthermore, participants were more accurate in Statistical compared to the Contextual association trials (*t*(83) = 5.224, *p* < .001, Cohen’s *d* = .472).

To test whether the Contextual associations resulted in explicit memory for the specific target location over and above just the side of the scene, we measured the Euclidian distance between the memory-recall response (Fig 3a) and the location of the target in the attention-orienting task. This target-response distance error value was then divided by the total possible distance error to generate the normalized localization error in percentage units (Fig 3c; see methods for further details). The location of the target in the orienting task provided a usable anchor to estimate error in the Statistical and Random conditions. For the Contextual condition, the location of the target in the attention-orienting task was the same as in the visual-search task. To ensure that localization was superior to just identifying the target side, only trials with correct hemifield responses were considered. Mean localization error was 16.85 (*SD* = 4.42) in the Contextual condition, 23.06 (*SD* = 4.01) in the Statistical condition, and 22.43 (*SD* = 4.00) in the Random condition. The ANOVA confirmed a main effect of Association type (*F*(2, 166) = 65.623, *p* <.001, *η2* = .314; Fig 3c). As predicted, participants were more precise in placing the remembered target in the Contextual associations than either the Random (*t*(83) = 9.230, *p* < .001, Cohen’s *d* = 1.324) or Statistical (*t*(83) = 10.095, *p* <.001, Cohen’s *d* = 1.472) conditions. There was no difference in precision between the Statistical and Random conditions (*t*(83) = 1.102, *p* = .863, Cohen’s *d* = .157). Indeed, when accounting for placing the target in the correct hemifield, performance in Statistical and Random association trials were not significantly different than chance (Statistical: *t*(83) = .622, *p* = .536, Cohen’s *d* = .068; Random: *t*(83) = .814, *p* = .418, Cohen’s *d* = .089).

## General discussion

We successfully introduced a task for investigating and comparing how different types of memory associations and regularities can improve performance in the context of a common visual-search task. By manipulating contextual associations vs. statistical regularities of target locations according to scene categories, we showed effective incidental learning of both types of memory relative to a random-association baseline condition over repeated visual-search blocks. Having demonstrated behavioural benefits from the incidental learning of these different types of association, we then showed that contextual vs. statistical regularities may impact attention-orienting with different temporal properties. A subsequent surprise explicit memory task confirmed that participants formed explicitly available memories regarding the target side for scenes with statistical regularities between scene category and target side as well as memories regarding scene-specific target locations for scenes with specific object-scene spatial contextual associations during visual search. Our findings highlight the promise of investigating learning and utilization of different types of memory associations within a common framework. They extend previous reports [46,53] by using natural scene stimuli to approximate everyday behaviour and by intermixing trials with different types of association within the same task to avoid confounds related to state variables.

We developed a task that promoted learning of two different types of associations within a common setting. Utilizing a single, visual-search task with embedded associations and regularities enabled equating stimulus materials, response requirements, and state variables between conditions. In the visual search task, independent of the type of association or regularity concerning target location, participants searched for a target within different scenes and reported its identity. They were not made aware of the experimental manipulation to association type. Including the random-association condition captured effects related to overall familiarity with the task and provided a continuous comparison point from which behavioural benefits in the other associations could be measured. Learning was measured over two successive days to allow time for incidental learning effects to emerge.

Despite the online nature of the visual-search task – participants showed good performance and learning of contextual associations and statistical regularities. Performance improvements occurred for all three conditions, displaying the importance of accounting for general learning related to task familiarity (see also 25). Specific learning and utilization of contextual and statistical associations were also noted by demonstrating benefits relative to the random condition. As we hypothesized, contextual associations brought greater benefits to visual search, with improved accuracy as well as response times over the two sessions. As predicted, contextual associations yielded a steeper learning rate compared to statistical regularities. Response times decreased more over blocks in the contextual-association condition compared to both other conditions. Response times were also faster in the statistical-association condition than the random-association condition on both days, but the decrease in response times over blocks did not differ significantly. The steeper benefit from contextual associations accords with previous studies manipulating contextual associations and statistical regularities in the form of probabilistic cueing in simple search arrays. However, the learning rates for statistical regularities was weaker than previously reported [53]. It is possible that the online testing added variance that precluded observing a reliable learning slope for the statistical condition. Alternatively, some of the choices in task parameters may have contributed, such as the use of natural scenes; adopting a deterministic, as opposed to probabilistic, statistical regularity; cueing side and not quadrant; and restricting the regularity to the spatial location of the target and not the response. Our challenging task, which required perceptual discrimination of the target, may have rendered response times less sensitive. The early benefits in accuracy for the contextual condition speaks to this form of association being particularly effective at changing perceptual sensitivity. Together, the pattern suggests the two types of memories may improve visual search performance through non-overlapping modulatory mechanisms. Future studies that systematically alter sensory and motor parameters within visual search should prove helpful in testing this possibility and providing more clues about the different modulatory pathways.

The different impact of the incidentally learned contextual and statistical associations on attention functions was corroborated by performance in the memory-guided attention-orienting task on the third day of testing. The memory-guided attention approach has the advantage of separating ongoing learning from testing the consequences of learning on attention. Within the visual-search context, many factors are at play when responding to one display, such as learning, finding the target, overcoming distraction, and preparing a response. Accordingly, it is difficult to isolate the effects of orienting attention. Including a separate task tuned for investigating spatial orienting after learning has stabilised lessens the impact of some of these other variables [45,47–49,64–66].

Our results showed a different effect of contextual associations and statistical regularities as a function of the interval between the scene, acting as the memory cue, and the scene array containing the target to be discriminated. Contextual associations significantly improved discrimination accuracy at the long interval, whereas the statistical regularity conferred a marginal benefit only at the short interval. The effects at the short interval were difficult to interpret with confidence, given the weak statistical result and the lack of difference between performance in the contextual and statistical conditions. It was not possible to conclude for certain if one memory condition influenced attention earlier, but it was clearer that the effects of contextual associations outlasted those of statistical regularities. An important caveat to the interpretations is that the overall memory strength of statistical regularities may have been weaker. Future studies may wish to bolster statistical learning before comparing the two types of memory head-to-head. It may be possible to strengthen the relative strength of the statistical regularities by making them more spatially focused (e.g., cueing a quadrant instead of a side) or making the search and orienting tasks less dependent on fine perceptual discriminations. Overall, we argue that using attention-orienting tasks may prove an effective way of charting the time course with which different types of memory associations prepare perceptual and motor functions. The approach complements studies comparing response times for recalling striatal-based and hippocampal-based memories [67].

The explicit memory task revealed that the nature of the associations learned incidentally during search were available for explicit report. Statistical regularities led to the best performance in remembering the side where targets had been learned. The improved identification of target side for the statistical-regularity condition may reflect the advantages of learning a compressed relationship between a scene category and a target side [19,21,25,68,69]. Contextual associations led to accurate reports of specific target locations learned within individual scenes over and above identifying the target side. Learning the location of target objects in individual scenes may provide effective contextual attentional templates that can improve performance across different tasks [64,70]. In our case, we observed how incidentally learned contextual associations improved visual search, perceptual discrimination in an attention-orienting task, and localization reports in explicit memory recall.

Together, these results provide a nice platform to investigate whether different neural systems that have been implicated in contextual or statistical learning can support performance benefits through different mechanisms and in different time frames. By varying the stimulus parameters and demands in the orienting task it may be possible to disentangle differences in the psychological consequences of contextual and statistical memories for guiding attention. For example, the tasks employed here stressed perceptual sensitivity and used two different intervals between the memory cue and the target array. Subsequent work could vary sensory and motor demands of the task as well as the intervals between cue and target scenes to reveal the time scales at which contextual and statistical memories can modulate sensory and motor stages of processing. Future implementations of this task could also examine whether the different memory systems interact competitively or synergistically during attentional guidance [71–73]. Brain-imaging and human neurophysiological studies will be useful for examining whether and how the different memory systems proactively modulate different stages of information processing.

Online testing brought advantages and limitations to data collection and interpretation. Advantages included facilitating data collection from a broader and more diverse participant pool as well as increased speed of data collection compared to in-lab testing. The disadvantages included the lack of control over the stimulus display and response equipment (e.g., screen size, distance from eyes to the screen, quality of mouse and keyboard, etc.), variability in timing responses (e.g., computer lag, network quality, etc.), and possible distractions during the task. However, decreased attention in online compared with in-lab testing is not a consistent finding [74,75], and online data quality is often comparable to in-lab studies [76,77]. Also, online testing limited the range of possible behavioural measures. For example, recording eye movements would provide rich additional information. Finally, the attrition rate, particularly for the multi-day study, was high compared to in-lab testing. Future studies should replicate this work in the lab as well as try hybrid approaches. One possibility is that a combination of learning occurring online and the final testing occurring in the lab would ensure strong learning followed by high levels of experimental control during the critical assessments.

Overall, we have provided new evidence that the nature of learned associations has important implications for their future utility in attention-orienting and memory-recall tasks. For example, we demonstrated that associations linked to functionally different memory systems can guide attention orienting, but that this guidance may unfold with different time courses. Benefits in explicit spatial memory recall also differed according to association type. These patterns of facilitation may suggest that there are functional differences in how the memory systems supporting contextual versus statistical learning benefit attentional orienting and memory recall. The behavioural effects established in this study provide a starting point for subsequent neuroscientific studies that investigate how these different memory systems support behaviour and how this memory-based regulation selectively breaks down in different neurodegenerative conditions. One exciting possibility is that follow-up studies could implement a similar design to measure disorders affecting the hippocampus [78–80] or the striatum [73,81,82] in a single task.

## Acknowledgements

We thank Melvin Kallmayer for his help with coding the original learning task.

